# Segmentation of Noisy Signals Generated By a Nanopore

**DOI:** 10.1101/014258

**Authors:** Jacob Schreiber, Kevin Karplus

**Affiliations:** Nanopore Group, Department of Biomolecular Engineering University of California, Santa Cruz; California, USA

**Keywords:** Sequence Analysis, Bayesian Statistics, Step Detection

## Abstract

Nanopore-based single-molecule sequencing techniques exploit ionic current steps produced as biomolecules pass through a pore to reconstruct properties of the sequence. A key task in analyzing complex nanopore data is discovering the boundaries between these steps, which has traditionally been done in research labs by hand. We present an automated method of analyzing nanopore data, by detecting regions of ionic current corresponding to the translocation of a biomolecule, and then segmenting the region. The segmenter uses a divide-and-conquer method to recursively discover boundary points, with an implementation that works several times faster than real time and that can handle low-pass filtered signals.

## INTRODUCTION

Nanopore devices consist of a tiny hole in an insulating barrier separating two wells containing salt solution, often a single protein porin inserted into a lipid bilayer (Kasianowicz *et al*, 1996). As a voltage is applied, ions pass through the nanopore, and ionic current readings are recorded by electrodes on either side of the bilayer. Biomolecules translocate through the nanopore, blocking the passage of ions and causing the ionic current to fluctuate in a sequence specific manner. The translocation of a biomolecule is usually mediated by an enzyme, which slows the passage of the biomolecule considerably, and gives it a stepwise quality (Cherf *et al*, 2012). This stepwise signal has a large noise added to it, which can be modeled as independent samples from a Gaussian distribution.

Two tasks that lend themselves to automation are the detection of translocation events and the segmentation of these events into steps that correspond to the stepwise movement of the biomolecule through the nanopore. Segmentation can be complicated by adjacent segments having similar ionic current, with the change being substantially less than the additive noise. A good segmentation technique must be able to distinguish these segments from each other, while not oversegmenting on noise.

The recorded ionic current from nanopore experiments is typically filtered to remove high-frequency noise before digitizing (Maitra *et al*, 2012; Nivala *et al*, 2013; Quanjun *et al*, 2012; Venta *et al*, 2013). When a signal is filtered, samples are replaced with samples that are influenced by the surrounding samples, making the noise no longer independent. Although we started with the simplifying assumption of independence, we have extended the method to handle signals filtered using a low-pass filter with an arbitrary cutoff frequency.

Currently, the analysis of nanopore data typically involves either summary statistics of a whole event (Manrao *et al*, 2011; Olasagasti *et al*, 2010; Sathe *et al*, 2011), such as duration and mean current, or segmentation performed by hand (Laszlo *et al*, 2013; Schreiber *et al*, 2013). Segmentation done by hand involves arbitrary decisions about whether a transition occurred when segmenting noisy signals and can introduce bias into the results. While it is true that the recent Oxford MinION device has a fully automated segmentation algorithm, it is proprietary and thus difficult to compare against. We introduce a simple rule-based parser to detect events and a recursive segmenter which takes into account low-pass filtration with an arbitrary cutoff frequency.

## 1 EVENT DETECTION

Data from nanopore experiments is recorded as a time series of ionic current. A simple model describes the data as coming from one of two states: *open-channel* where nothing is passing through the nanopore, and the ionic current levels are high, or *event*, where something is passing through the nanopore, and the ionic current levels are lowered.

The sharp transitions between the two states makes a rule-based parser appropriate, where an event is identified as a region of ionic current below a threshold. An example of an ionic current trace is shown in Fig. 1, with a red bar showing the threshold of 90 pA used.

**Figure 1.**
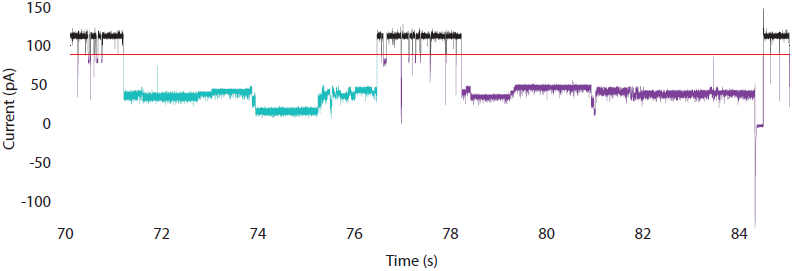
The event detector was run on a sample region of ionic current. The red line indicates the threshold at 90 pA used in this case. The purple events have been rejected as either being too short (duration less than 1 second) or having a negative minimum ionic current (manually clearing a blocked pore). The teal event is the only event accepted by the event detector.

Simple thresholding results in many uninformative events, falling into two classes: *transient events*, corresponding to the translocation of molecules without an enzyme slowing them down; and *blockage* events, where the movement of the biomolecule through the nanopore has stalled, and the pore was cleared by reversing the voltage bias. Transient events can be identified by their very short duration, and blockage events by the transient negative ionic current that clears the pore. A minimum duration threshold and a minimum current threshold are used to remove the uninformative events. While many events are initially identified in Fig. 1a and colored purple, only the single teal-colored event is accepted.

## 2 SEGMENTATION

### 2.1 Sample Scoring

While some nanopore analyses focus on summary statistics (such as mean current, duration, or rms noise) of a complete translocation event, others require summary statistics for each of the individual ionic current segments which make up the event. To compute these statistics, the boundaries between adjacent segments must be identified. An automated segmentation method must be able to identify these boundaries even when adjacent segments are not very different.

Two popular groups of boundary-finding algorithms are filter-derivative methods (Basseville, 1981) and Bayesian methods (Girshick & Rubin, 1952), both of which are described in a textbook on detecting abrupt changes (Basseville-Nikiforov, 1993)—our proposed segmenter is a Bayesian method. Boundary points are identified by our segmenter one at a time using a recursive algorithm. We start by considering the entire event as a single segment, then consider each possible boundary to break it into two segments. To avoid edge effects and ensure that all segments have at least a minimum duration, only potential boundaries more than the minimum segment length from the ends of the segment are considered.

Each possible boundary is scored using a log-likelihood function (Eq. 1). If the maximal score is above a threshold, the segment is split and the two subsegments are recursively segmented. The recursion terminates either when the segment to split is less than twice the minimum segment length or no score within the segment exceeds the threshold.

The score function

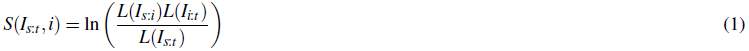

is the log-likelihood ratio of two models applied to a region of ionic current *I*_*s*:*t*_: a two-segment model split at boundary *i*, and a single-segment model. Each segment is modeled as an independent, identically distributed process with a Gaussian distribution having parameters *μ* and *σ* selected to maximize the likelihood of the corresponding segment. The likelihood is simply the product of the probabilities of each sample in the region given the model:

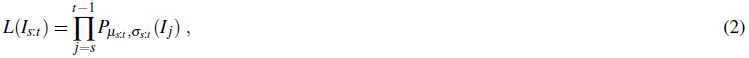

and the probability of a single sample given a Gaussian distribution with parameters *μ* and *σ* is

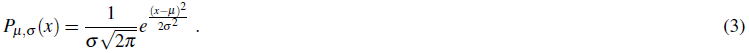

Combining Eq. 3 and Eq. 2 and taking the log, we get Eq. 4 for the log-likelihood of a region of ionic current:

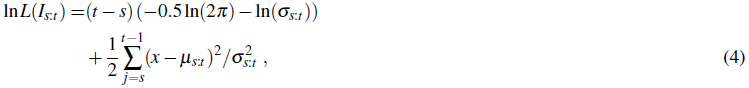

Because we chose *μ*_*s*:*t*_ and *σ*_*s*:*t*_ using maximum-likelihood estimates, the sum simplifies to *t* – *s*, giving us

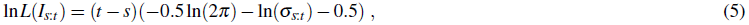

which depends only on *σ*_*s*:*t*_, not on *μ*_*s*:*t*_. Using Eq. 5, we can simplify the score in Eq. 1 to

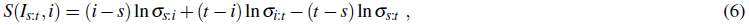

which will always be non-negative.

### 2.2 Score Threshold for *i.i.d*. Samples

At the end of each recursion step, the segmenter has calculated the log-odds score for each possible boundary in a region of ionic current. The score of the maximally scoring sample is then compared to a threshold, and if it is above the threshold, the boundary is accepted. For a full Bayesian treatment, we want to use the posterior log-odds, not just log-likelihood.

The posterior-odds ratio is the likelihood ratio multiplied by the prior-odds ratio, which can be parameterized by the expected number of segments per second (sps), and the sampling frequency of the discrete time steps in the data (*f*_*s*_). The prior probability of a sample being a boundary is 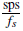, and the prior odds ratio is 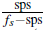. Using the log-likelihood ratio in Eq. 6, the log of the posterior-odds ratio is then *S*(*I*_*s*:*t*_, *i*) + ln(sps) – ln(*f*_*s*_ – sps), and we choose boundaries where the maximal value of the score has

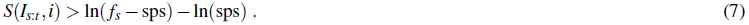

For those uncomfortable with Bayesian statistics, we can also use a frequentist approach to set a threshold based on the p-value of the scores, so that we can estimate and control the expected number of false positives. Unfortunately, we do not have a theoretically proven model for how *S*(*I*_*s*:*t*_, *i*) is distributed, so we did a number of simulations of Gaussian i.i.d. sequences and found empirically that the score appears to follow a simple exponential distribution:

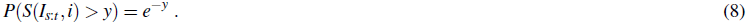

This distribution is independent of the length of the window being scored or the parameters *μ* and *σ* of the underlying Gaussian distribution, though there may be some edge effects for samples very close to existing boundaries (perhaps because we are using population estimates of *σ* rather than unbiased estimates). Even the independence of the distribution with respect to length is not quite right—short windows have slightly higher scores in them than long ones do, though still exponentially distributed (*e*^−*λy*^ for *λ* < 1). The effect is fairly small for the windows we are interested in, and so we have not attempted to characterize it precisely, but treat all scores as coming from a simple exponential distribution.

This assumption of simple exponential distribution of scores allows us to estimate the number of false positive boundary calls to expect and set a threshold to control the number. If we set the acceptable number of false positive boundary calls per second (FPS), then we want the probability of this score arising by chance for any single sample to be less than 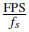. The maximal log score across a region of unfiltered ionic current must satisfy

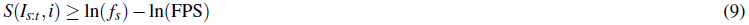

for us to accept splitting the region.

Subsequent steps in the analysis pipeline may be better able to handle extra boundaries than missing boundaries (or vice versa), so controlling the types of error that the segmenter makes can be useful, whether by changing the prior (sps) or acceptable false positives per second (FPS).

The analysis in this section can be extended to any modeling scheme that results in i.i.d. Gaussian residuals. For example, we have used it to segment signals into stepwise sloping segments (instead of stepwise constant segments) using linear regression for each segment.

The segmentation method also works well when the segments are distinguished by changes in the amount of noise, even when the mean values remain the same, unlike filtered-derivative approaches to segmentation, which attempt to remove the noise before segmenting.

### 2.3 Score Threshold for Filtered Samples

Section 2.2 gave thresholds for data with i.i.d noise, but most nanopore-generated data is filtered to remove transient spikes and high-frequency noise before digitizing. When a low-pass filter is applied to the data, the samples are no longer independent, as each point has been replaced by a point influenced by its neighbors. The scores increase with low-pass filtering, as nearby data values are closer to each other, so dividing up a segment provides a larger decrease in standard deviation.

Fig. 2a shows the distribution of scores for Gaussian noise at a sampling frequency of 100kHz, low-pass filtered at 10kHz, 5kHz, 2kHz, and 1kHz. The exponential fit, assuming a cutoff frequency of *rf*_*s*_/2 (that is, *r* times the Nyquist frequency) is *P*(*S*(*x*_*s*:*t*_, *i*) > *y*) = *e*^−*rx*^.

Thus for low-pass filtered data with cutoff *rf*_*s*_/2, we want

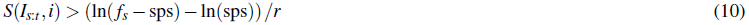

or

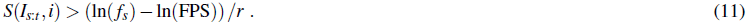

**Figure 2.**
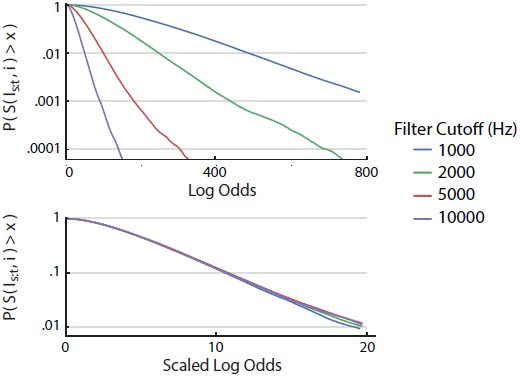
Cumulative distribution functions (CDFs) of the score produced by running the segmenter on 1000 regions each with 10000 randomly generated samples at 100kHz from a single Gaussian distribution, filtered with various low-pass 5th-order Bessel filters. (a) The CDF appears to be exponential when the signal is filtered with a low-pass Bessel filter, but with increased scores the lower the filter cutoff used. (b) A simple scaling by the ratio between the cutoff frequency and the sampling frequency makes all the CDFs essentially the same. A slight divergence is noted at high scores, due to the small number of samples having rare values.

### 2.4 Examples

A randomly generated signal comprised of four segments was generated with 100kHz sampling for illustrating segmentation. This signal was filtered using a first-order low-pass Bessel filter with a cutoff of 5kHz, as shown in Fig. 3a. The segmenter was then run on this region of data, and produced the correct segmentation indicated by the vertical red lines.

We calculated the threshold from Eq. 10 with *f*_*s*_ = 100 kHz, sps = 1 segments per second, and 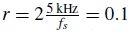, to get a threshold of 115.1 for a boundary to be called. The segmenter scored every sample in the region according to Eq. 6 (Fig. 3b). Since the maximally scoring sample scored above threshold, it was taken as a boundary. The process was repeated on the region to the left of this boundary (Fig. 3c), where another boundary is found. No boundaries were found in the region to the left (Fig. 3d) or right (Fig. 3e) of that boundary, causing the region to the right of the first boundary to be scanned (Fig. 3f). A boundary was found, but no further boundaries were found on either the left (Fig. 3g) or the right (Fig. 3h) of that boundary. The process then terminated, with the fully segmented event.

**Figure 3.**
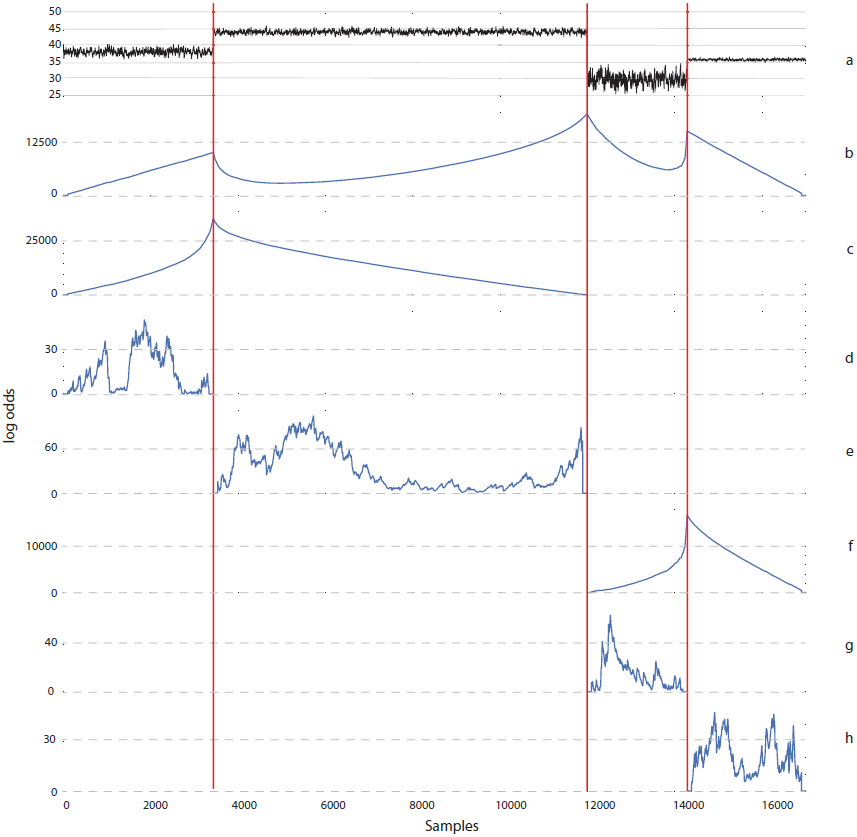
An example event being segmented. The filtered event (a) is scored at every time step using the log-odds of a boundary being called there. The maximally scoring time step is taken as a boundary. The region to the left of that point is then analyzed in the same manner (b), producing another boundary. The regions to the left (d) and right (e) of that point are analyzed, but produce low scores, and so no further boundaries are found. The region to the right of the original boundary is then analyzed, with a single boundary being found (f), but no further on either side (g&h) of that.

To illustrate the effect of the sps parameter on real data, the segmenter was run with a range of priors on 127 events collected from a DNA nanopore experiment. There is an increase in the number of observed segments as the prior increases. The plot is divided into two main regions: on the left where the likelihood score dominates over the prior when calculating the posterior and the number of observed segments is relatively constant, and on the right where the prior dominates over the likelihood score and the number of observed segments increases proportional to the prior. When the prior approaches the sampling frequency, the segmenter begins to call segments averaging about 1.3 times the minimum duration parameter. With a minimum duration of 100 samples and a sampling frequency of 100kHz the observed number of segments will plateau around 800 segments/second (Fig. 5).

**Figure 4.**
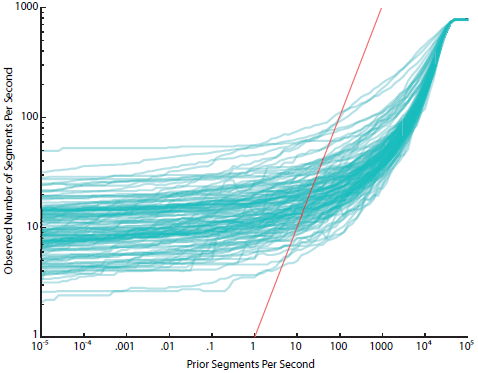
Number of segments per event for differing prior numbers of segments per second, for 100kHz data low-pass filtered to 2kHz. At low priors, the likelihood score overwhelms the prior, causing a relatively constant number of segments, while at high priors the prior overwhelms the likelihood score. The minimum segment duration parameter (here, 100 samples) causes a plateau when the prior nears the sampling frequency, due to early termination of the recursion. This early termination is also responsible for the number of observed segments per second being much lower than the prior, even when the prior is dominating the data. A x=y line is shown in red to show the difference caused by the early termination.

**Figure 5.**
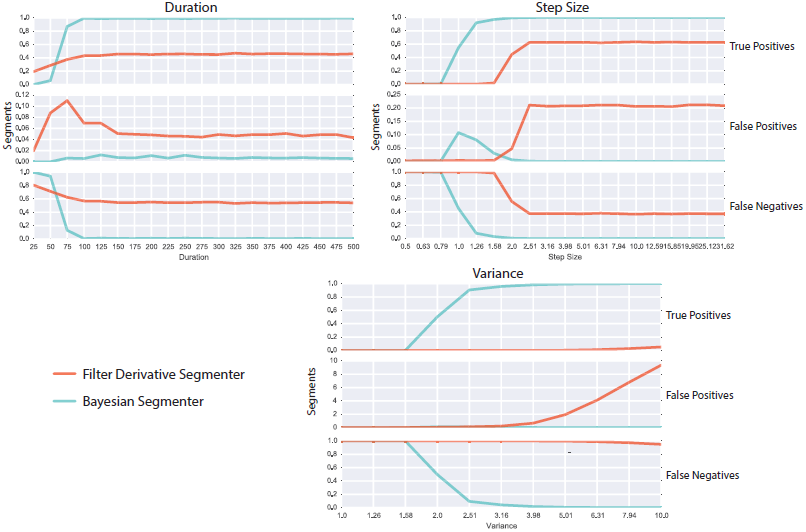
Evaluation of the filter-derivative segmenter versus our segmenter. Our segmenter required less tuning and performed better in difficult cases.

### 2.5 Comparison to Filter-Derivative Segmenters

In order to evaluate this segmenter, we ran it against a hand-tuned filter-derivative segmenter on three major tasks; (1) detecting a single step of two picoamps while varying the duration in samples of the first segment, (2) detecting a single step of varying step size in two equal duration segments, and (3) detecting a ‘step’ between two segments with equal mean but varying variance. We measured the average number of true positive, false positive, and false negative segmentations across 100,000 random events generated with the same parameters. We said that a segmentation was correct if the transition was detected within five samples of the correct transition.

When the duration of the first segment is varied, the Bayesian segmenter picks up on the change very early on, getting a single correct

## 3 IMPLEMENTATION

This process for detecting and segmenting events in nanopore signals should run in real time; either segmenting a stream of data as it comes in or quickly segmenting an event shortly after its completion. To test the speed of the algorithm, the event detector was implemented in Python and the segmenter was implemented in Cython, a language that allows the efficiency of C within Python. The current implementation is designed to segment full events and is available at the first author’s public github page.

To measure the speed of the implementation, example data was collected from an experiment 37.5 minutes long, with a sampling frequency of 100 kHz, filtered before digitizing using a 5 kHz, 4-pole, low-pass Bessel filter. The event detector was run on three segments of data, each 12.5 minutes long, and 127 events were detected. The segmenter was then run on one event at a time. The whole process, including loading the data, detecting events, and segmenting events, took approximately 88 seconds to process 2250 seconds of data on a Intel Core i5-3470 CPU clocked at 3.2GHz. The segmenter by itself took 71 seconds to segment the 903 seconds of data identified as events (about 0.5 seconds per event), making it a little over 12 times faster than real time.

The current implementation allows for the segmenter to look only at short windows (typically 1 or 2 seconds), which would permit segmentation to occur in slightly delayed real-time, if the I/O for the program were changed. When no boundary is found in a window, the window is moved forward by half its length and the process repeated. If a window is split, the right hand portion is extended to a full window width, to avoid artifacts from the artificial window boundary.

Both Bayesian and frequentist thresholds are supported by the software, using (ln(*f*_*s*_ – sps) – ln(sps) + ln(*f*_*s*_) – ln(FPS)) /*r* as the threshold, and defaulting sps = *f*_*s*_/2 and FPS = *f*_*s*_ so that the unused parameter is ignored.

In order to show that the implementation worked, the segmenter was run on a filtered event using a prior sps value of 40, the approximate catalytic rate of (Φ29 (Lieberman *et al*, 2010; Soengas *et al*, 1995), and a cutoff frequency of 2kHz (Fig. 6). 58 total segments were detected on an event lasting 67 seconds, giving an observed rate of 52 segments per second. The boundaries called by the segmenter are very close to those which would be called by hand. This example trace shows an example of many adjacent segments with similar means but different variances being correctly identified, a key advantage of this algorithm, being underlined in cyan. Also shown are examples of short, distinct segments being identified, such as the one underlined with a red line, but other short spikes not being called segments in noisier signal, such as the one underlined with the orange line.

**Figure 6.**
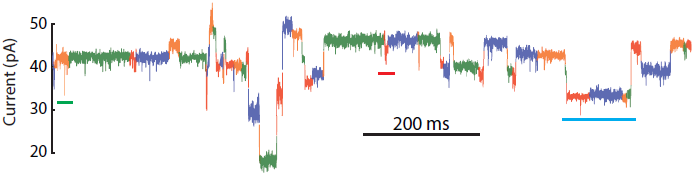
An example event segmented with a prior segments per second of 40 after filtering using a low-pass Bessel filter with a cutoff frequency of 2kHz. Segments are colored in a red-blue-orange-green cycle, where boundaries are indicated by a change in color. Three important phenomena are marked with colored lines: in green, short spikes in somewhat noisy signal are not called as segments; in red, short but distinct segments still are identified; and in teal, several adjacent segments with similar means but differing variances can still be distinguished.

## 4 CONCLUSION

The automatic analysis of nanopore signal starts with event detection and segmentation of these events into useful subunits. Event detection can easily be achieved using a rule-based filter given the large, abrupt drops in ionic current as a biomolecule passes through a nanopore, but segmenting these events is more challenging. We presented a recursive segmenter, with both a Bayesian and a frequentist interpretation of the data, which can handle low-pass filtered data with an arbitrary cutoff frequency.

An implementation of this method has been shown to work significantly faster than real time.

Future work on the software includes implementing the algorithm on a computer directly connected to the digitizer for the nanopore signal, so that real-time analysis of the data can be used to control the nanopore (such as detecting blockages and automatically reversing the current to clear the pore).

## ACKNOWLEDGEMENT

The authors would like to acknowledge Zachary Wescoe, who ran the nanopore experiments used as an example in Figures 1, 5, and 6.

KK was responsible for the initial development and analysis of the segementation algorithm. JS was responsible for the event detector, the Cython implementation, the release in the PyPore package, and the figures in this article.

## Funding

This work was partially supported by the National Human Genome Research Institute (grant HG006321-02)

## REFERENCES

Basseville,M. (1981) Edge detection using sequential methods for change in level—Part II: Sequential detection of change in mean, IEEE Trans. Acoustics, Speech, Signal Processing, 29, 32–50.

Basseville,M. and Nikiforov,I. (1993) Detection of Abrupt Changes: Theory and Application Prentice Hall. ftp://ftp.irisa.fr/local/as/mb/k11.pdf

Cherf, G. et al. (2012) Automated forward and reverse ratcheting of DNA in a nanopore at 5-Å precision, Nature Biotechnology, 30, 344–348.

Girshick,M.A. and Rubin,H. (1952) A Bayes approach to a quality control model, Annals Mathematical Statistics, 23, 114–125.

Kasianowicz,J. et al. (1996) Characterization of individual polynucleotide molecules using a membrane channel, Proc Natl Acad Sci, 93(24), 13770–13773.

Laszlo,A. et al. (2013) Detection and mapping of 5-methylcytosine and 5-hydroxymethylcytosine with nanopore MspA, Proc Natl Acad Sci, 110(47), 18904–18909.

Lieberman,K. et al. (2010) Processive Replication of Single DNA Molecules in a Nanopore Catalyzed by phi29 DNA Polymerase, J. Am. Chem. Soc., 132(50), 17961–17972.

Maitra,R. et al. (2012) Recent advances in nanopore sequencing, Electrophoresis, 33(23), 3418–3428.

Manrao,E et al. (2011) Nucleotide Discrimination with DNA Immobilized in the MspA Nanopore, PLoS One, 6(10), e25723.

Nivala,J. et al. (2013) Unfoldase-mediated protein translocation through an *α*-hemolysin nanopore, Nat Biotechnol, 31(3), 247–250.

Olasagasti,F. et al. (2010) Replication of Individual DNA Molecules under Electronic Control Using a Protein Nanopore, Nat Nanotechnol, 5(11), 798–806.

Quanjun,L. et al. (2012) Voltage-Driven Translocation of DNA through a High Throughput Conical Solid-State Nanopore, PLoS One, 7(9), e46014

Sathe,C. et al. (2011) Computational Investigation of DNA Detection using Graphene Nanopores, ACS Nano, 5(11), 8842–8851.

Schreiber,J. et al. (2013) Error Rates for Nanopore Discrimination Among Cytosine, Methylcytosine, and Hydroxymethylcytosine Along Individual DNA Strands, Proc Natl Acad Sci, 110(47), 18910–18915.

Soengas,M. et al. (1995) Helix-destabilizing Activity of Φ29 Single-stranded DNA Binding Protein: Effect on the Elongation Rate During Strand Displacement DNA Replication, J. Mol. Biol., 253, 517–529.

Venta,K. et al. (2013) Differentiation of Short Single-Stranded DNA Homopolymers in Solid-State Nanopores, ACS Nano, 7(5), 4629–4636.

